# Phase Separation of Shell Protein and Enzyme: An Insight into the Biogenesis of a Prokaryotic Metabolosome

**DOI:** 10.1101/2022.01.14.476392

**Authors:** Gaurav Kumar, Sharmistha Sinha

**Affiliations:** Chemical Biology Unit, Institute of Nano Science and Technology, Knowledge city, Sector-81, Mohali-140306, India

**Keywords:** Bacterial Microcompartments, Shell Proteins, Phase Separation, Molecular Crowding

## Abstract

Bacterial microcompartments are substrate specific metabolic modules that are conditionally expressed in certain bacterial species. These all protein structures have size in the range of 100-150 nm and are formed by the self-assembly of thousands of protein subunits, all encoded by genes belonging to a single operon. The operon contains genes that encode for both enzymes and shell proteins. The shell proteins self-assemble to form the outer coat of the compartment and enzymes are encapsulated within. A perplexing question in MCP biology is to understand the mechanism which governs the formation of these small yet complex assemblages of proteins. In this work we use 1,2-propanediol utilization microcompartments (PduMCP) as a paradigm to identify the factors that drive the self-assembly of MCP proteins. We find that a major shell protein PduBB’ tend to self-assemble under macromolecular crowded environment and suitable ionic strength. Microscopic visualization and biophysical studies reveal phase separation to be the principle mechanism behind the self-association of shell protein in the presence of salts and macromolecular crowding. The shell protein PduBB’ interacts with the enzyme diol-dehydratase PduCDE and co-assemble into phase separated liquid droplets. The co-assembly of PduCDE and PduBB’ results in the enhancement of catalytic activity of the enzyme. A combination of spectroscopic and biochemical techniques shows the relevance of divalent cation Mg^2+^ in providing stability to intact PduMCP *in vivo*. Together our results suggest a combination of protein-protein interactions and phase separation guiding the self-assembly of Pdu shell protein and enzyme in solution phase.

**Significance Statement:** Present work shows how surrounding environment modulates the self-assembly behavior of a major shell protein of 1,2-propanediol utilization microcompartment (PduMCP). Appropriate ionic strength and macromolecular crowding bring about liquid-liquid phase separation of the shell protein. Under crowded environment Mg^2+^ displayed unique property to drive the formation of shell protein liquid condensates. The co-phase separation of a native enzyme along with the shell protein enables it to outperform the enzyme in isolation. This adds on to the existing concept of phase separation being the underlying principle behind the genesis of lipid free organelles in prokaryotes. Further, our results indicate that the divalent metal ion Mg^2+^ plays an intricate role in the outfitting of the structure-function integrity of PduMCPs.

**Graphical Abstract:** 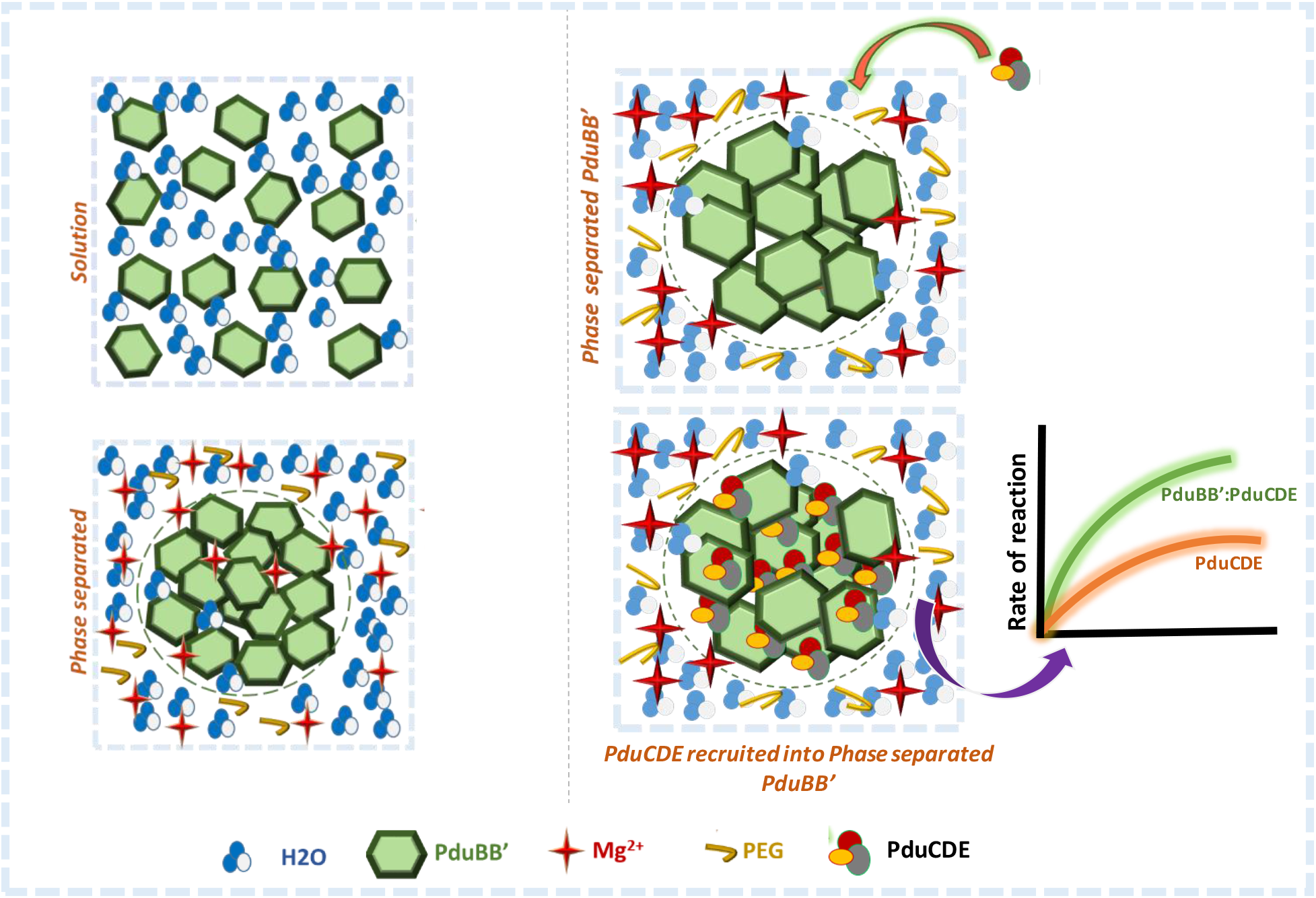

## Introduction

Living cells synergize multiple biochemical pathways by organizing cellular reactions within sub-cellular organelles.^1,2^ Based on the composition of the outer cover of these organelles, they are categorized as membrane-bound and membrane-less organelles. Membrane bound organelles such as mitochondria and Golgi bodies have outer coat made of lipid bilayer, whereas membrane less organelles like nucleolus, cajal bodies and p-granules lack lipid bilayer outer membrane and are solely made of proteins.^3–6^ Membrane less organelles are not exclusive to eukaryotic cells, but have been reported in prokaryotes in the form of phase separated protein condensates such as Ftz-SlmA complex or ParABS system.^7,8^ Many bacterial species have also evolved to make use of nano-sized proteinaceous microcompartments for their energy metabolism that help them to survive under energy deficient conditions. Bacterial Microcompartments (MCPs) are a unique embodiment of an organelle that house a congregation of enzymes bound by a distinct outer cover made of shell proteins. The outer shell unlike typical membranes lack the lipid component and is made up of entirely proteins. The shell proteins act as physical barrier regulating the movement of biomolecules from cytoplasm to the lumen of MCPs and vice versa.^9,10^

Over the years MCPs have been extensively studied for their role in bacteria survival, pathogenesis and their potential in the development of synthetic nano-reactors.^11–13^ It is therefore important to understand how multiple subunits of proteins are driven towards one another and orchestrated into small yet complex catalytic reactors. In recent past, the concept of phase separation has been put forward in the context of membraneless organelle formation, including the biogenesis of MCPs such as carboxysomes and pyrenoids^14,15–18^. The ability of shell protein to undergo liquid-liquid phase separation has been proposed to be crucial step in the genesis of these MCPs. This motivates us to question if phase separation is indeed a driving force behind the formation of PduMCPs. Since PduMCPs are a complex assortment of multiple number of proteins, studying the self-assembly properties of individual PduMCP proteins might be a pragmatic approach towards deciphering the assembly principles of whole PduMCP. Our initial bioinformatics study using catGranule server ^19^ suggests that out of the 8 shell proteins of PduMCP, the major shell protein PduBB’ has the highest propensity to undergo LLPS **(Fig. 1a, i)**.

**Figure. 1:**
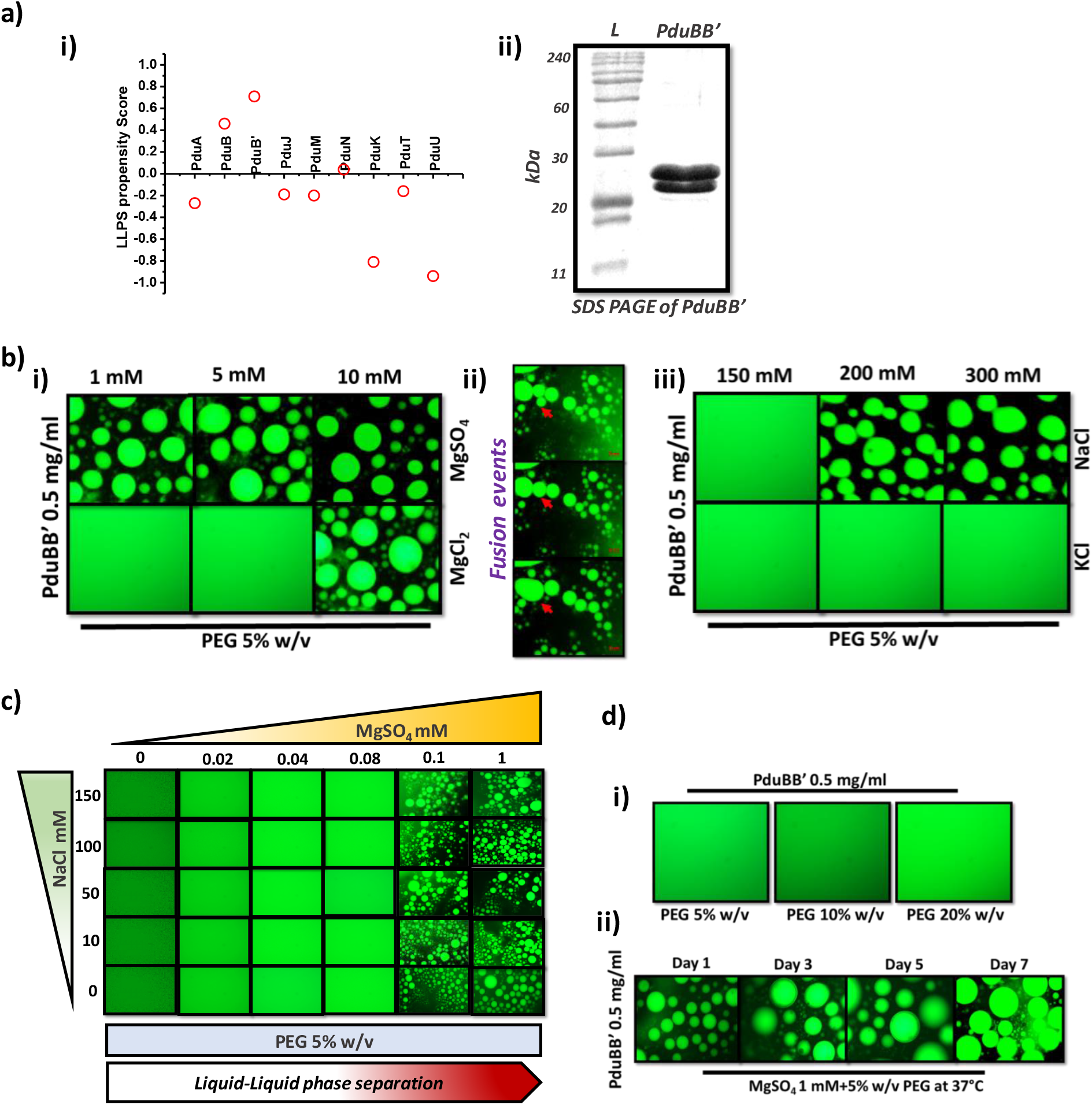
LLPS of shell protein PduBB’. a, (i) Liquid-liquid phase separation propensity of Pdu shell proteins predicted using catGranule webserver19, (ii) SDS PAGE of PduBB’, b) Microscopic visualization of self-assembly dynamics of PduBB’ in solution phase in the presence of salts and crowding agent (PEG-8000, 5% w/v): (i) Liquid-liquid phase separation of PduBB’ in the presence of MgSO_4_ (at 1 mM, 5 mM and 10 mM concentrations), and MgCl_2_ (at 10 mM concentration), (ii) Fusion of protein droplets confirming liquid nature of the PduBB’ droplets, (iii) Liquid-liquid phase separation of PduBB’ in the presence of NaCl (at 200 mM and 300 mM concentrations), c) Phase diagram showing phase separation of PduBB’ in solution phase at different concentrations of NaCl and MgSO_4_, d) i) Microscopic visualization of Alexa-488 labeled PduBB’ in in the presence of 5% w/v-20% w/v PEG-6000, ii) Day dependent microscopic visualization of PduBB’ droplets after incubating the protein samples at 37°C.

The trimeric shell protein PduBB’ is an essential component of PduMCP and is required for the formation of PduMCP^20,21^. It is expressed as a combination of two constituent proteins, PduB (270 aa) and PduB’ (233 aa) **(Fig. 1a, ii)**. Absence of this shell protein impairs the formation of PduMCP in *Salmonella enterica*,^20^ highlighting its significance in the PduMCP biogenesis. The composition of the component proteins (PduB and PduB’) in a PduBB’ trimer is not known. The presence of extended disordered N-terminal region in PduB **(Fig. S1a)**, would also make it difficult to crystalize the full length PduB in PduBB’ protein. However, the available crystal structure of a homologous protein, PduB from *Lactobacillus reuteri*^22^ suggests the involvement of ionic interaction in the lateral associations of the trimeric shell protein assemblies **(Fig. S1b)**. Notably, the involvement of electrostatic interactions is a key factor in promoting higher order assembly of various shell proteins ^22–26^. The presence of disordered region and ionic interactions along with high LLPS propensity **(Fig. 1a, i)**, makes shell protein PduBB’ an ideal candidate for our study. Existing reports suggest that increased ionic strength of solution tends to reduce the LLPS of disordered proteins ^27 28^. Since PduBB’ has both structured and disordered domains, it would be interesting to check the effect of salts on the phase separation behavior of PduBB’ in solution phase. In the present study we use a series of spectroscopic and biophysical techniques to study the self-assembly of shell protein PduBB’and its co-assembly with the signature enzyme PduCDE. The consequences of the co-assembly on the catalytic efficiency of the enzyme is explored. Our results accentuate the significance Mg^2+^ ions in driving the phase separation of PduBB’ and in preserving the integrity of the intact PduMCP. Further, the difference in the phase separation behavior of PduBB’ and its truncated version PduB underscores the role of the extended N-terminal region in regulating the self-assembly of the shell protein in solution phase.

## Results

### Phase separation of shell protein PduBB’ in a crowded environment

The cellular microenvironment remains highly viscous and crowded due to the presence of many bio-macromolecules^29–31^. It is therefore important that we study the self-assembly dynamics of proteins under crowded environment. Polyethylene glycol (PEG) is a well-known crowding agent which has been extensively used for mimicking the cellular microenvironment *in vitro*.^32,33^ In order to visualize the dynamics of PduBB’ under crowded environment and at different ionic strengths, fluorescence microscopy of Alexa-488 labelled PduBB’ solution is performed in the presence of salts and PEG-6000 (5% w/v). We select Mg^2+^ salts (namely MgCl_2_ and MgSO_4_), that are important buffer ingredients needed for production as well as purification of PduMCP from *Salmonella enterica* 23. Distinct PduBB’ assemblies are seen in the presence Mg^2+^ salts **(Fig. 1b, i)**. Careful observation of the dynamics of the PduBB’ assemblies in real time, reveal their liquid like behavior, undergoing fusion in solution phase **(Fig. 1b, ii and Video Sv1)**. We observe that MgSO_4_ and MgCl_2_ are able to trigger the LLPS of PduBB’ at concentrations 1 mM and 10 mM respectively **(Fig. 1b, i)**. In comparison to Mg^2+^ salts, monovalent salt, namely NaCl induce LLPS of PduBB’ at high concentration of NaCl (200 mM and above) **(Fig. 1b, iii)**. The LLPS of PduBB’ is not observed in the presence of KCl under the conditions studied **(Fig. 1b, iii)**. The potential of the salts to trigger LLPS of PduBB’ follows the following order: MgSO_4_ >MgCl_2_ >NaCl > KCl.

We further see that that MgSO_4_ (0.1 mM) is sufficient to trigger the LLPS of shell protein PduBB’ and the presence of NaCl is not essential for the same **(Fig. 1c, Fig. S2)**. In the absence of any salt, the crowding agent PEG alone fails to trigger the LLPS of PduBB’ in solution **(Fig. 1d)**, indicating the importance of both ionic strength and macromolecular crowding in the formation shell protein liquid droplets. These liquid phases are found to be stable at 37°C for several days without any aggregation **(Fig. 1d)**. The LLPS of PduBB’ protein with increase in solution ionic strength suggests a role of hydrophobic interactions in bringing the shell protein molecules together. This is very likely, as shell proteins do not have buried hydrophobic core and have exposed hydrophobic patches on its surface^34^. Therefore, the presence of kosmotropic ions like Mg^2+^ would expel out water molecules from the surrounding environment of PduBB’, bringing the shell protein molecules closer due to hydrophobic interactions. Since Na^+^ ions are weak kosmotrope, a high concentration of NaCl is required to trigger LLPS of PduBB’.

We find that in the absence of any crowding agent salts tend to increase the size of PduBB’ assemblies **(Fig. S3)** and optical density of PduBB’ solution **(Fig. 2a)**. To decipher the underlying mechanism behind this observation, we take surface charge of PduBB’ into consideration. PduBB’ (with a PI value of 5.11) has an overall negative surface charge giving zeta potential value of −34.2 mV. Presence of salts increase the zeta potential value of PduBB’ solution indicating surface charge masking of the shell protein by metal ions present in the vicinity of PduBB’. This would reduce the electrostatic repulsion among the shell proteins inducing their self-association. Stronger surface charge masking of PduBB’ by Mg^2+^ ions explains why Mg^2+^ salts trigger the self-association of PduBB’ at very low concentration of 1 mM, indicating Mg^2+^ binding sites on the protein. Using the MIB webserver (http://bioinfo.cmu.edu.tw/MIB/)^35^, we theoretically predict the probable Mg2+ biding residues on the surface of PduBB’ **(Fig. S4)**. The resultant PduBB’ assemblies are seen as solid associates under the microscope. To check if these assemblies are soluble associates or irreversible aggregates, the protein samples are dialyzed in 10 mM phosphate buffer (pH 7.4) to remove the salts. Post dialysis, the PduBB’ solution becomes clear and we do not see any assembly under microscope **(Fig. 2b)**. Lifetime measurements of Alexa-488 labelled PduBB’ in the presence of salts showed no substantial difference indicating retention of structure upon salt induced self-assembly **(Fig. 2c and S5)**. Circular dichroism studies of salt treated PduBB’ samples post dialysis in 10 mM phosphate buffer (pH 7.4) show the preservation of the secondary structure **(Fig. 2d)**. These studies confirm that salts do not alter the secondary structure of the shell protein PduBB’ during its self-association in solution. Further, a combination of lifetime measurement studies, and fluorescence anisotropy experiments show that during the process of phase separation, the backbone of the PduBB’ protein remains solvated but the dynamics of the background remain restricted **(Supplementary Information, Fig. S6)**. Solvation of protein backbone would maintain the protein molecules in liquid phase and prevent aggregation. The restricted backbone flexibility may occur due to the increased solution viscosity as a result of macromolecular crowding.

**Figure 2:**
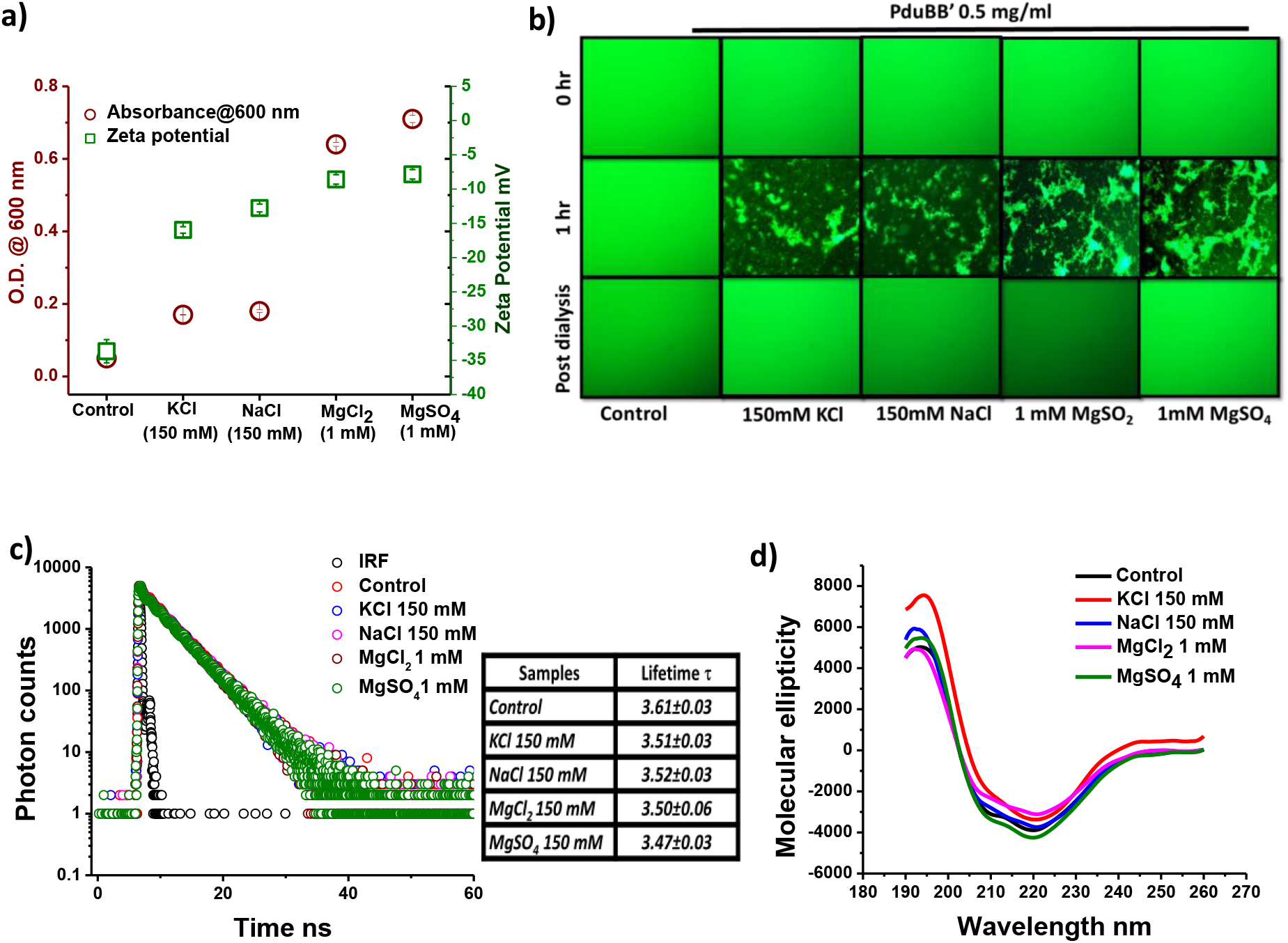
Mechanism behind salt driven self-assembly of PduBB’. a) Absorbance at 600 nm and zeta potential of PduBB’ solution in the absence and presence of salts, b) Microscopic visualization of assemblies of Alexa-488 labeled PduBB’ (0.5 mg/ml) in solution phase in the presence of salts. PduBB’ tend to self-associate within an hour forming visible soluble associates which disappear upon dialysis and removal of salts, c) Fluorescence lifetime of Alexa-488 labeled PduBB’ (0.5 mg/ml) in the absence and presence of salts, d) CD spectroscopy of PduBB’ (0.5 mg/ml) in the absence and presence of salts post dialysis.

### Shell protein has higher tendency than enzyme to undergo phase separation

The enzyme PduCDE is expressed a s a heterotrimer **(Fig. S7a)**. For comparing the LLPS behavior PduCDE and PduBB’, we look for their minimum concentrations needed to observe LLPS in solution phase. The LLPS of PduBB’ is observed up to very low shell protein concentration of 0.025 mg/ml while the enzyme undergoes phase separation only at a concentration of 1 mg/ml **(Fig. 3a and 3b)** suggesting that shell protein PduBB’ has a higher phase separation propensity than the enzyme PduCDE **(Fig. S7b)**. The understanding of the difference in the surface morphology of PduBB’ and PduCDE may explain the phase separation behavior of the two proteins. Unlike enzyme PduCDE which is a globular protein with buried hydrophobic residues, shell protein PduBB’ has exposed hydrophobic patches on its surface^34^. This is also evident from higher ANS fluorescence intensity of PduBB’ than that of PduCDE at same protein concentration (**Fig. 3c)**. The expulsion of water molecules by a kosmotropic salt MgSO_4_, would strongly trigger the self-association in case of PduBB’ with exposed hydrophobic patches. Bio-layer interferometry studies also show that PduBB’: PduBB’ interaction is stronger than PduCDE: PduCDE interaction **(Fig. 3d and 3e)**. For the PduBB’: PduBB’ interaction, no dissociation is observed post association event, indicating very strong association of the molecules **(Fig. 3d)**. A global fitting of the association and dissociation kinetics of PduCDE: PduCDE interaction gives a dissociation constant (*k_d_*) of 3.5 X 10^−6^ M **(Fig. 3e)**. This strong self-assembly of the shell proteins may be contributing factor to their higher tendency to undergo self-association and phase separation.

**Figure 3:**
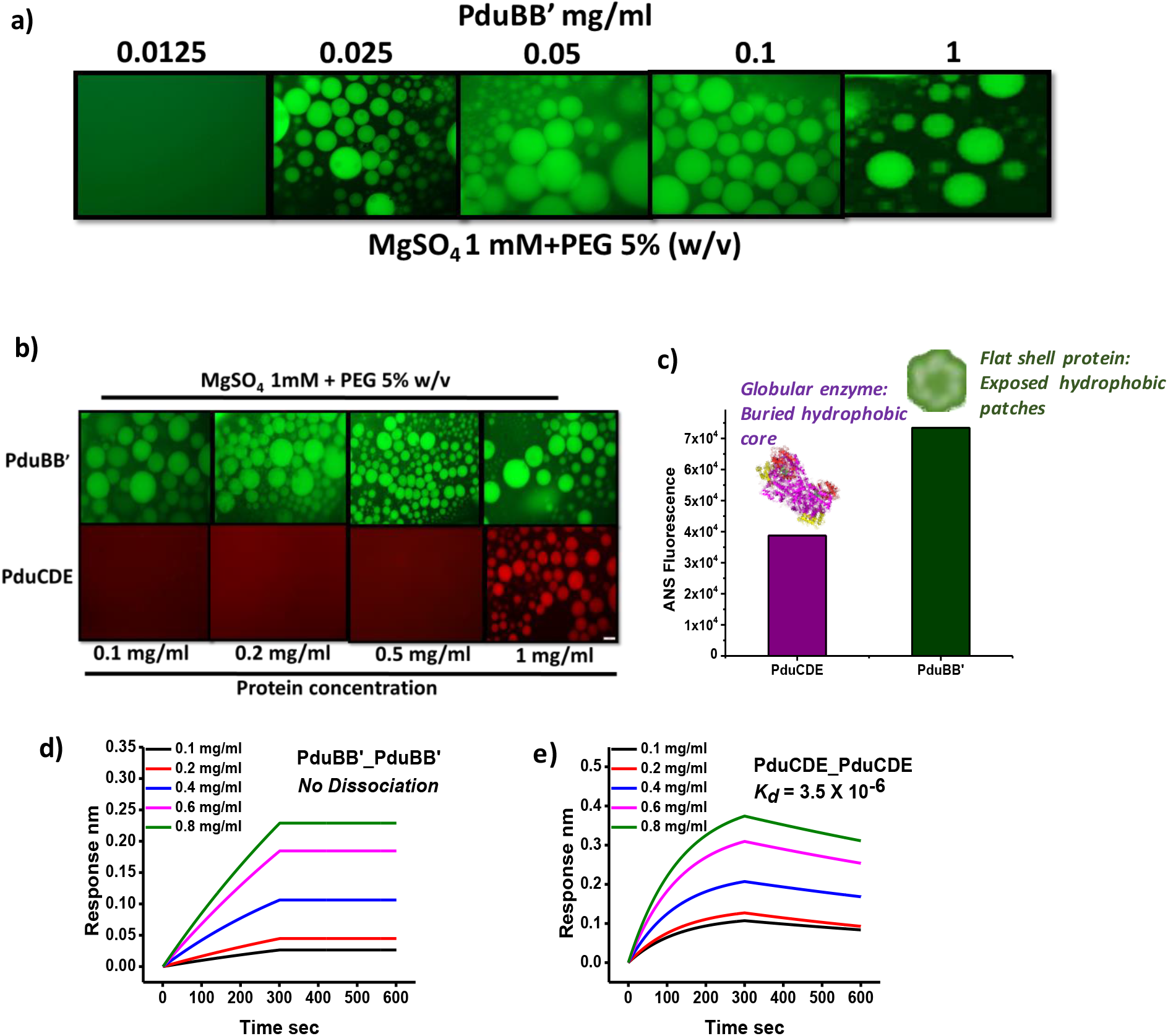
Shell protein PduBB’ has higher LLPS propensity than enzyme PduCDE. a) Liquid-liquid phase separation of PduBB’ at different protein concentration, b) Liquid-liquid phase separation of PduBB’ and PduCDE at protein concentrations 0.1, 0.2, 0.5 and 1 mg/ml, c) ANS fluorescence of PduCDE and PduBB’ at 0.5 mg/ml protein concentration, d) Protein-protein interaction performed using biolayer interferometry to show association and dissociation kinetics for PduBB’: PduBB’ association, e) Protein-protein interaction performed using biolayer interferometry to show association and dissociation kinetics for PduCDE: PduCDE interaction.

### Co-phase separation of PduBB’ and PduCDE

Our interferometry result suggests that a moderate to strong affinity exists between the shell protein PduBB’ and enzyme PduCDE with a dissociation constant (k_d_) of 2.7 X 10^−7^ **(Fig. S7c)**. In this section we study the phase separation behavior of PduBB’ and PduCDE in the presence of one another. In the presence of macromolecular crowding (5% PEG-6000) and 1mM MgSO_4_, PduBB’ and PduCDE, co-localize into protein droplets **(Fig. 4a)**. PduCDE co-separated with PduBB’ in LLPS shows enhanced enzyme activity compared to PduCDE alone as LLPS **(Fig. 4b)**. This result highlights the importance of both phase separation and shell protein: enzyme interaction in improving the catalytic activity of the enzyme PduCDE. We also observe that PduCDE could be recruited within pre-formed phase separated PduBB’ droplets **(Fig. 4c)**. PduCDE also gets recruited within phase separated PduBB’ solid associates (at 2 mg/ml) within 3 min of interaction. In soluble form, without any macromolecular crowding and salt the enzyme PduCDE shows a specific activity of 23.3±1.4 μmol min^−1^ mg^−1^. We look for the specific activity of PduCDE at different concentrations (0.5-2 mg/ml) under phase separated condition, in the absence and presence of PduBB’ **(Fig. 4d and 4e)**. In the absence of PduBB’, the enzyme PduCDE shows an activity of ~ 25.4±0.05 μmol min^−1^ mg^−1^ at protein concentration of 0.5 mg/ml **(Fig. 4d)**. An increment in specific activity of PduCDEis seen only at enzyme concentrations 1 mg/ml and above. This may be attributed to the fact that at a concentration below 1 mg/ml, PduCDE fails to undergo phase separation. Hence, at 0.5 mg/ml it exhibits an activity similar to the one seen in the absence of salt and crowding agent. However, In the presence of PduBB’ under similar phase separated condition, PduCDE shows significantly higher specific activity, highlighting the significance shell protein-enzyme interaction in modulating catalytic activity **(Fig. 4e)**. Further, the specific activity is enhanced by ~3 fold when the enzyme is recruited within solid phase at 2 mg/ml concentration of PduBB’ **(Fig. 4e)**. In the following sections, we attempt to understand importance of salts in driving the self-assembly of shell protein PduBB’ in solution.

**Figure 4:**
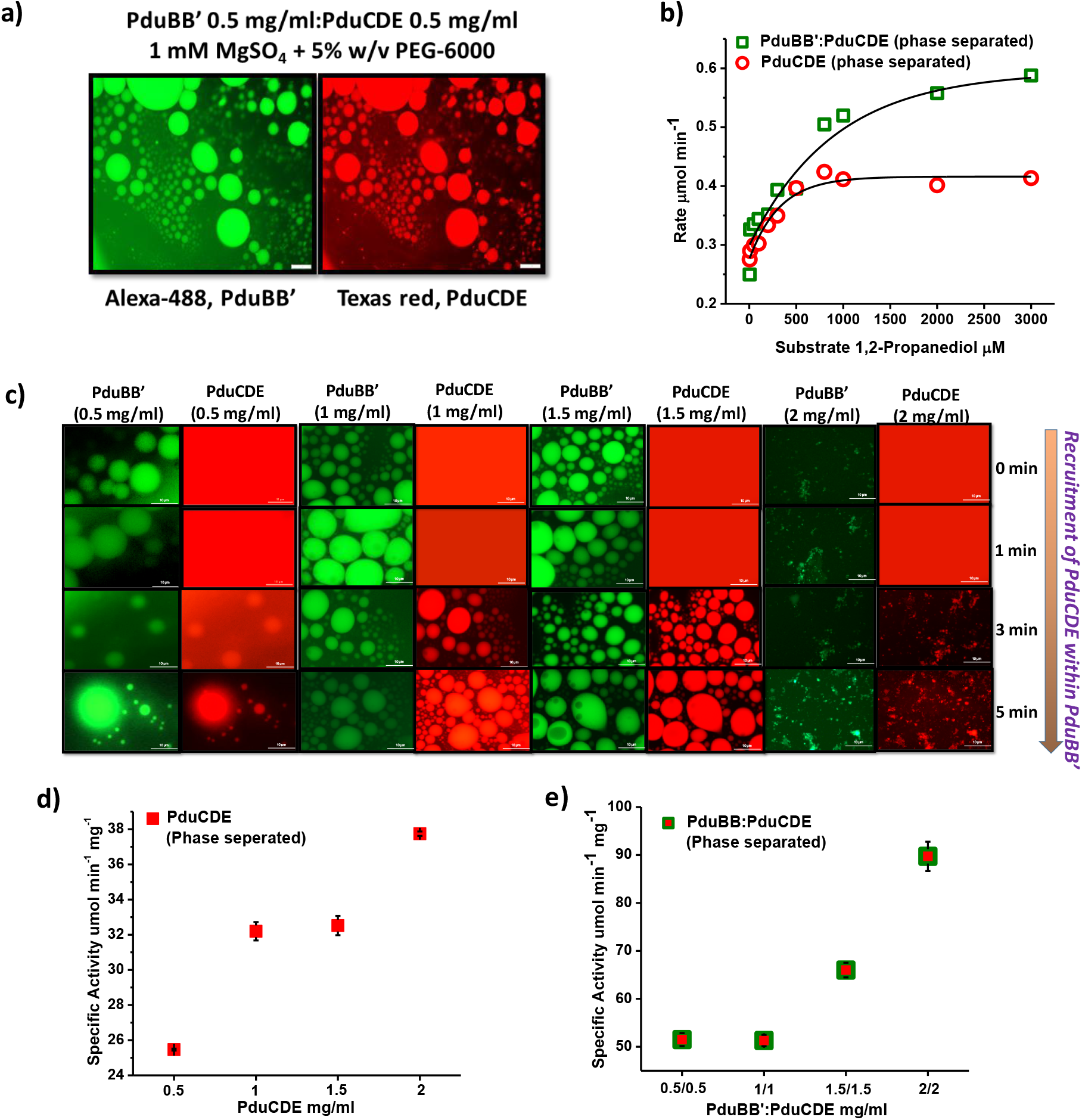
Co-assembly and phase separation of PduBB’ and PduCDE. a) Co-phase separation of PduBB’ and PduCDE, b) Diol dehydratase assay for PduCDE (1 mg/ml) and PduCDE:PduBB’ (1 mg/ml: 1 mg/ml) under phase separated condition at different substrate (1,2-propanediol) concentrations, c) Time dependent recruitment of Texas red-labelled PduCDE within Alexa-488 labelled phase separated PduBB’, d) Specific activity of PduCDE at different protein concentrations in the presence of 1 mM MgSO_4_ and 5%w/v PEG-6000, e) Specific activity of PduCDE at different shell protein: enzyme ratios in the presence of 1 mM MgSO_4_ and 5%w/v PEG-6000.

### Role of divalent metal ion in the structure-function relationship of Pdu microcompartment in vivo

The self-association and phase separation of the major shell protein PduBB’ in the presence of salts leads us to hypothesize that the ionic strength of environment may have crucial implications on the assembly, stability and functioning of intact PduMCP. To test this hypothesis, the diol dehydratase activity of purified PduMCPs is determined in the presence and absence of salts in its storage buffer (in 50 mM Tris buffer (pH 8.0) containing 50 mM KCl and 5 mM MgCl_2_) **(Fig. 5a)**.. The salts are removed by dialyzing the PduMCPs against Tris buffer (pH 8.0) without salts. Upon removal of salts from their surrounding environment, the purified PduMCPs exhibit lower catalytic activity, **(Fig. 5a)**. This suggests that the presence of salts is necessary for the optimum catalytic activity of PduMCPs. Next, we assess how removal of only Mg^2+^ affects the stability and optimal functioning of PduMCP. For this experiment, purified PduMCP is incubated in the presence of EDTA (0 mM, 2 mM, 5 mM and 30 mM) for 5 min, followed by intrinsic fluorescence measurement at temperatures ranging from 20°C-90°C. **Fig. 5b** shows derivative plot of intrinsic fluorescence of PduMCP as a function of temperature, which gives us the melting temperature (T*_m_*) of PduMCP. The increase in EDTA concentration results in the decrease in the T*_m_* of PduMCP, suggesting lower stability of PduMCP in the presence of EDTA. The chelation of Mg^2+^ by EDTA also lowers the catalytic activity of the purified PduMCP. **(Fig. 5c)**. It is however important to note that while 5 mM EDTA results in ~24% decrease in PduMCP activity, no further significant reduction in PduMCP activity is observed upon increasing the EDTA concentration to 30 mM. This indicates that the chelation of Mg^2+^ ions by EDTA may only destabilize the outer shell of PduMCP, without resulting in its complete breakdown. Size determination of the PduMCPs in the presence and absence of EDTA show no significant change in the hydrodynamic diameter of PduMCP, confirming that PduMCP remains intact upon chelation of Mg^2+^ by EDTA (**Fig. 5d)**. Since the presence of Mg^2+^ ions drives the self-association of shell protein, its removal would reduce the stability of the outer shell of PduMCP, lowering its catalytic efficiency and thermal stability. This idea is strengthened by the observation where the chelation of Mg^2+^ by EDTA impairs the LLPS of PduBB’: PduCDE mixture **(Fig. S8a)**, suggesting that the absence of Mg^2+^ significantly affects the self-assembly behavior of Pdu proteins in solution. Together these results highlight the role of Mg^2+^ ions in maintaining stability and optimum level of activity in intact PduMCP. The presence of Mg2+ ion itself does not influence the catalytic activity of the enzyme PduCDE alone (**Fig. S8b)**.

**Figure 5:**
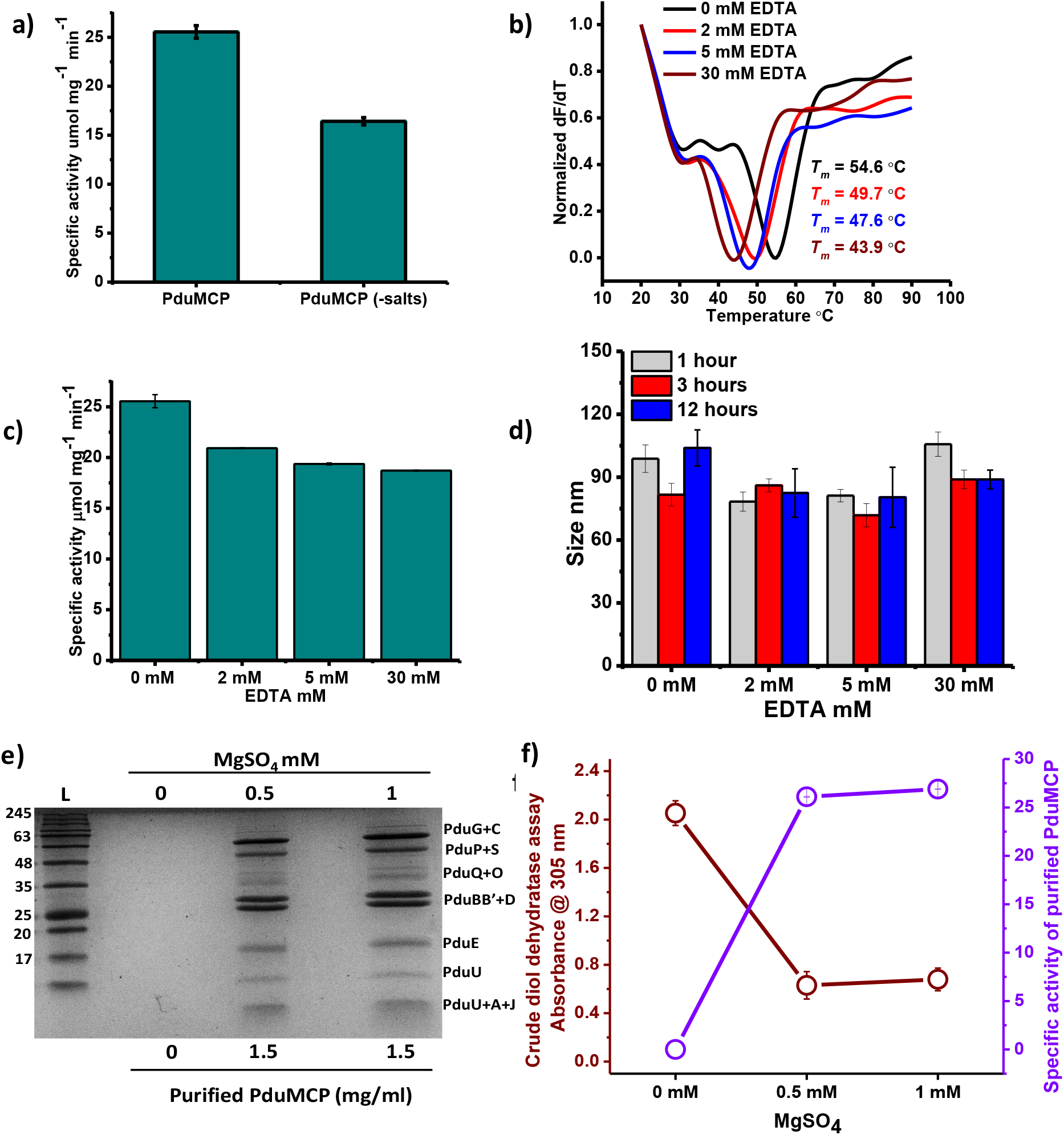
Effect of Mg^2+^ ions on PduMCP assembly. a) Diol dehydratse activity of PduMCP in the presence and absence of salts (50 mM KCl and 5 mM MgCl_2_), b) Derivative plot of intrinsic fluorescence of PduMCP in the absence and presence of EDTA, c) Diol dehydratase assay of PduMCP in the absence and presence of EDTA, d) Hydrodynamic diameter of PduMCP in the absence and post treatment of EDTA for 1 h, 3 h and 12 h, e) SDS PAGE of PduMCP purified from *Salmonella enterica LT2* grown in the absence and presence of 0.5 mM and 1 mM MgSO_4_, f) Crude diol dehydratase assay using cell lysate of Salmonella LT2 grown in the absence and presence of 0.5 mM and 1 mM MgSO_4_ (shown in maroon), Specific activity of purified PduMCP from Salmonella LT2 grown in the absence and presence of 0.5 mM and 1 mM MgSO_4_ (shown in purple).

Motivated by these observations, we next question the importance of Mg^2+^ ions in formation of a functional PduMCP *in vivo* by inducing PduMCP formation in the presence and absence of Mg^2+^ ions. Surprisingly, in the absence MgSO_4_, PduMCPs failed to show any protein bands in SDS PAGE gel **(Fig. 5e)**. On the other hand, PduMCPs are successfully purified from culture grown in the presence of MgSO_4_ (0.5 mM and 1 mM), and exhibit optimum diol dehydratase activity **(Fig. 5f)**. Interestingly, diol dehydratase assay performed using crude cell lysate of *Salmonella enterica* suggests that Pdu component proteins are generated in all cultures irrespective of MgSO_4_ concentration **(Fig. 5f)**. Notably, the production of propionaldehyde is higher in case of *Salmonella enterica* grown in the absence of MgSO_4_. Higher propionaldehyde production suggests that in the absence of MgSO_4_, the outer shell of PduMCP is comparatively more porous, exposing the encapsulated enzymes^23,36^. This may be attributed to the fact that MgSO_4_ strongly triggers the self-assembly of the major shell protein in solution phase. Besides, we have also observed that the enzyme PduCDE, in the presence of PduBB’ and 1mM MgSO_4_, undergoes co-separation and co-condensation in solution **(Supplementary information Fig. S9)**. Therefore, the absence of Mg^2+^ salt from the growth medium may hinder the formation of an intact PduMCP and render the outer shell of the PduMCP more porous. This experiment shows that the presence of MgSO_4_ in the growth medium is essential for the production of a fully intact PduMCP. In the following section we attempt to get an insight into the physical mechanism behind the salt driven self-assembly of shell protein in a cell, by mimicking a crowded cellular microenvironment *in vitro*.

## Discussion

Understanding the self-assembly mechanism of a multi-component system like bacterial microcompartment remains a challenge. Their complex organization calls for a minimalistic approach, where the factors controlling the self-assembly of their individual components are identified in isolation. The factors identified could be then applied towards understanding the functioning of the entire microcompartments. In this regard, we attempt to probe the self-assembly behavior of a major shell protein (PduBB’) and the signature enzyme (PduCDE) of PduMCP in solution phase. We begin our study by looking at the effect of ionic strength of and macromolecular crowding of the surrounding environment on PduBB’ assembly. We observe that the shell protein PduBB’ takes the shape of spherical droplets displaying liquid like behavior. This is an interesting observation as in recent years, phase separation has been suggested to be the principle mechanism governing the self-assembly of many different proteins at sub-cellular level^37^. The de-mixing of protein molecules in a crowded cellular microenvironment not only drives protein aggregation^28,38,39^ but also results in the formation of membraneless organelles^40,41^. The concept of phase separation has been recently floated in the context of carboxysome biogenesis^42,43^. Elegantly designed microscopic studies indicate the ability of carboxysome shell protein to undergo phase separation. These reports along with our results suggest that the mechanism of phase separation may play a crucial role in the biogenesis of several MCPs including PduMCP.

The combination of surface charge masking and kosmotropic effect of metal ions appear to be the underlying cause of salt driven phase separation of shell protein PduBB’. In the absence of crowding agent, the shell protein PduBB’ tends to self-associate in solution phase without undergoing any significant change in its secondary structure. The effect of salts on the self-assembly of shell protein has been reported earlier, where the hexameric shell protein from *Haliungium Ocraceum* was found to self-associate with increase in ionic strength.^44^. The effect of divalent metal ions was found to be significantly higher than that of monovalent metal ions, an observation similar to the one made in the present study. We believe that ions present in the cellular environment may play crucial role in modulating the self-assembly of proteins of many different MCPs. It is therefore important that beside protein-protein interaction, the solution ionic strength is taken into consideration while discussing the self-assembly of various MCP proteins. Our *in vitro* data on phase separation of PduBB’ points out at the significance of Mg^2+^ ions on shell protein assembly, and is reflected in both *in vitro* and *in vivo* studies using PduMCP.

PduBB’ form *Salmonella enterica* is composed of two proteins, encoded by two overlapping genes^21^. To date, this unique combination of shell proteins has been reported exclusively for PduMCP. The physiological relevance of this quirky combination is an interesting subject of study. Our *in vitro* work has shown that while PduBB’ has the potential to undergo LLPS in solution phase, the smaller component of the shell protein (PduB’, 233 aa) forms solid associates under similar environmental condition **(Fig. S10)**. It is likely that the presence of the disordered N-terminal in PduB component in PduBB’ helps to regulate the self-assembly of this shell protein, highlighting the importance of the naturally tuned combination of two shell proteins in PduMCP. Further *in vivo* cellular studies would throw more light on this subject. This venture would require the use of high resolution imaging techniques that have recently gained attention for visualizing the phase separation of proteins in bacterial cells^8^.

## Methods

### Expression and Purification of PduBB’ and PduCDE

Transformed *E.coli* BL21 cells (transformed with genes for PduCDE or shell protein cloned into pTA925 vector, containing Kanamycin resistant gene) are selected on kanamycin plate. A single colony is selected and inoculated in 10 ml of LB broth medium containing 50 μg/ml of kanamycin and grown overnight at 37°C. For the expression of PduCDE, 1% of overnight grown primary culture of *E.coli* BL21DE3 (transformed with genes for PduCDE) is inoculated in 400 ml of LB media (containing 50 μg/ml kanamycin) and incubated at 37°C for 1.5 to 2 h until the cells grow up to an OD 600 of 0.5. The expression of PduCDE is induced by adding 1 mM IPTG and incubating the culture at 37°C for 4 h. For the xpression of PduBB’, 1% of overnight grown primary culture of *E.coli* BL21DE3 (transformed with genes for PduBB’) is inoculated in 400 ml of LB media (containing 50 μg/ml kanamycin) and incubated at 37°C for 1.5 to 2 h until the cells grow up to an OD 600 of 0.5. The expression of PduBB’ is induced by adding 0.5 mM IPTG and incubating the culture for 12 h at 28°C. The cells are then harvested and lysed in column buffer (50 mM Tris-base pH 7.5, 200 mM NaCl and 5 mM imidazole). The supernatant is passed through Ni-NTA column. The non-specific proteins are removed by washing the column with wash buffer (50 mM Tris-base pH 7.5, 200 mM NaCl, 50 mM imidazole) and protein of interest is eluted by passing elution buffer (50 mM Tris-base pH 7.5, 200 mM NaCl, 200 mM imidazole). The eluted protein samples are dialyzed in 10 mM sodium phosphate buffer (pH 7.4) to remove NaCl and imidazole. Protein concentration is estimated using Bradford’s reagent and the purity of the proteins is checked by running SDS PAGE.

### Purification of PduMCP

Purification of PduMCP for *Salmonella enterica LT2* is done as described earlier^45^. Briefly, 1% of overnight grown culture of *Salmonella enterica LT2* is inoculated in 400 ml minimal media (1X NCE, non-carbon E), supplemented with 0.6% 1,2-PD, 0.5% succinic acid and 1 mM MgSO_4_ (For the experiment shown in **Fig. 5e and 5f**, the MgSO_4_ concentration is varied from 0 mM to 1mM). It is incubated at 37°C for 16h. The harvested cells are washed with buffer A (50 mM Tris Base pH 8, 500 mM KCl, 25 mM NaCl, 12.5 mM MgCl_2_, 1.5 % 1,2-PD) at 8000 X g for 5 min at 4 °C. The washed cells are re-suspended in Buffer A containing 75% bacterial protein extraction reagent (BPER-II), 1 mg/ml of lysozyme, 2 mg DNase and 0.4 mM phenyl methane sulfonyl fluoride (PMSF). The cells are then kept on a shaker at 45-50 rpm at room temperature for 30 min, followed by incubation on ice for 5 min. The lysed cell debris are removed by centrifugation (12,000 X g for 5 min, 4 °C) and PduMCPs in the supernatant are pelleted by centrifugation (20000 X g for 20 min, 4 °C). The PduMCP pellet is re-suspended in Buffer A containing 60% of B-PER II and 0.4 mM PMSF and is centrifuged at 20,000 X g for 20 min at 4°C. The supernatant is discarded and the thin film of PduMCP obtained is re-suspended in pre-chilled Buffer B (50 mM Tris Base pH 8, 50 mM KCl, 5 mM MgCl_2_, 1 % 1,2-PD). The re-suspended thin film is centrifuged at 12,000 X g for 5 min at 4 °C. The supernatant containing PduMCP is collected and stored at 4°C for further use. The concentration of PduMCP is estimated using Bradford’s reagent and purity is checked using SDS PAGE.

### Dynamic Light scattering and Zeta potential

To study the effect of salts on the size distribution of shell protein PduBB’, the shell protein samples (0.1 mg/ml) are incubated in the presence of different concentrations of salts (KCl, NaCl, MgCl_2_, and MgSO_4_) for 1h at room temperature. The size distribution of the shell protein is recorded using ZetaSizer Nano ZSP (Malvern Instruments, UK). The scattered intensity at backscattered angle 173° is recorded, and for each sample 3 recordings are taken. The intensity percentage for each recording is plotted at different concentrations of salts. To get the maximum population of shell protein assemblies in solution, number percentage of each reading is noted and their mean and standard error are plotted at different concentrations of salts. To determine the effect of salts on the surface charge of PduBB’, 200 μl of PduBB’ solution is incubated in the presence of different salts (KCl, NaCl, MgCl_2_, and MgSO_4_) for 1 h at room temperature. Zeta potential of each sample is measured using ZetaSizer Nano ZSP (Malvern Instruments, UK).

### Turbidity Assay

200 μl of shell protein PduBB’ samples (0.5 mg/ml) are incubated in the absence and presence of salts (KCl, NaCl, MgCl_2_, and MgSO_4_) at room temperature for 1 h. The turbidity of the protein samples are measured in 96 well plate using multiplate reader Infinite M Plex (Tecan, Austria). To check the effect of MgSO_4_ on the self-association of enzyme PduCDE, 200 μl of PduCDE sample is incubated in the absence and presence of 1 mM MgSO_4_. The turbidity of the PduCDE samples are measured in 96 well plate using multiplate reader Infinite M Plex (Tecan, Austria).

### Circular Dichroism

PduBB’ sample incubated in the presence of salts are dialyzed against phosphate buffer (pH 7.4). The dialyzed samples are used for determination of secondary structure of PduBB’ protein using a CD spectrophotometer (Jasco J-1500, CD spectrophotometer, (Jasco, Japan). For the determination of CD spectra 200 μl of samples are taken in a quartz cell of path length of (0.1 cm) in a N_2_ atmosphere. CD spectra are measured in the far-UV region (195-260 nm). For each sample, three scans with scanning speed of 200 nm/min is accumulated. Sample holder temperature is maintained at 25 °C using a mini circulation water bath (Jasco MCB-100) connected to the water-jacketed sample holder chamber.

### Fluorescence lifetime measurements

Alexa-488 labeled PduBB’ samples are incubated in the presence of different salts (KCl, NaCl, MgCl_2_, and MgSO_4_) for 1 h at room temperature. Fluorescence lifetime of the samples is determined using TCSPC system, HORIBA DeltaFlex (Horiba, Japan). The samples are excited using 402 laser diode and emission is recorded at 520 nm at bandwidth of 2 nm. Before recording the lifetime decay of the samples the instrument response factor is determined using Ludox solution. The life time of the samples is calculated by fitting the initial 10 ns of the decay curve using mono-exponential decay equation.

### Fluorescence microscopy and visualization of phase separation

For visualization of PduBB’ associates, Alex-488 labeled PduBB’ samples are incubated in the presence of salts for 1h at room temperature. 20 μl of protein samples are drop casted onto a glass slide and visualized under the FITC channel using upright fluorescence microscope, model: OLYMPUS BX53 (Olympus, USA). To check the effect of macromolecular crowding on the self-assembly of PduBB’ or PduCDE, 2 μl of PEG-6000 (50% w/v stock) is added to 20 μl of PduBB’ or PduCDE samples in the presence of salts. The samples are drop casted onto a glass slide and visualized under the FITC channel using upright fluorescence microscope, model: OLYMPUS BX53 (Olympus, USA). For co-phase separation experiment, 2 μl of PEG-6000 (50% w/v stock) is added to 20 μl of protein samples (contacting a mixture of Alexa-488 labeled PduBB’ and Texas red labeled PduCDE) in the presence of 1 mM MgSO_4_. The samples are drop casted onto a glass slide and visualized under the FITC and Texas Red channels using upright fluorescence microscope, model: OLYMPUS BX53 (Olympus, USA).

### Fluorescence anisotropy measurements

Alexa-488 labeled PduBB’ is mixed with 5 % w/v of PEG-6000 and 1mM MgSO_4_ to induce phase separation. Time resolved fluorescence anisotropy of the PduBB’ samples under control and phase separated condition is determined using TCSPC system, HORIBA DeltaFlex (Horiba, Japan). Rotational correlation time is determined by fitting the anisotropy decay curve to mono exponential decay equation.

### Biolayer interferometry

Protein-protein interaction studies are carried out using Fortebio OctateK2 (Molecular Devices, USA). 0.5 mg/ml of PduBB’ or PduCDE (PduCDE in case of enzyme-shell interaction study) is loaded onto activated Amine reactive second generation (AR2G) biosensors. Prior to loading, the AR2G biosensors are activated using a solution containing 40mM 1-ethyl-3-(3-dimethylaminopropyl) carbodiimide (EDC) and 20 mM of N-hydroxysuccinimide (NHS) mixed in 2:1 molar ratio. The protein loaded sensors are then immersed in ethanolamine (pH 8.0) to block the sites on sensor where protein molecules are not bound. Then the sensors are immersed in titer wells containing protein samples at different concentrations to record their association and dissociation kinetics. The dissociation constant is obtained by global fitting of the association and dissociation kinetics is performed using Data Analysis HT 9.0.0.33 software provided with the OctateK2 instrument.

### Diol-dehydratase assay

Diol dehydratase activity of purified PduMCP or PduCDE is estimated by 3-methyl-2-benzothiazoline hydrazine (MBTH) method. For routine assays, 2 μg of enzyme (for PduCDE assay) and or 5 μg of PduMCP (for PduMCP assay) is added to 900 μl of the assay buffer (0.2 M 1,2-propanediol, 0.05 M KCl, 0.035 M potassium phosphate buffer (pH 8.0) at 37 °C. For the assay of phase separated samples shown in **Fig. 4b**, reaction mixture contains 1 mg/ml of PduCDE in control sample and 1 mg/ml of both shell protein and enzyme in sample containing the PduBB’ and PduCDE mixture. In the experiment shown in **Fig. 4c**, the concentrations mentioned in the figure represent the concentrations of PduBB’ and PduCDE in the reacrtion mixture.

Reaction is started by adding 50 μl of adenosyl cobalamin AdoCbl (15 μM) and quenched after 10 min by adding 1 ml of potassium citrate buffer (pH 3.6). Then, 0.5 ml of 0.1 % MBTH (w/v) is added to the reaction mixture and the it is incubated at 37 °C for 15 min. After 15 min, 1 ml of dH_2_O is added and absorbance of the product formed is taken at 305 nm using UV spectrophotometer. Absorbance at 304 nm is converted to the concentration of the product formed using molar extinction coefficient 13000 cm^−1^. We define specific activity as μmol of product formed by 1 mg of PduCDE or PduMCP in 1 min.

To assess the effect of EDTA on the activity of PduMCP, the purified PduMCP samples (0.1 mg/ml) are incubated in the absence and presence of different concentrations of EDTA 5 min at room temperature. The EDTA treated and control samples are then used for diol dehydratase assay.

For the crude cell lysate assay shown in **Fig. 5f**, 3 ml of *Salmonella enterica* cells in the growth medium are centrifuged (post O.D 600 normalization) to get the pellet. The pellets are dissolved in 500 μl of Buffer A 75% bacterial protein extraction reagent (BPER-II). The cells are dispersed and kept at room temperature for 30 min for lysis. Post lysis, the cell debris is removed and 50 μl of the supernatant is used for the diol dehydratase assay as described above.

### Temperature dependent fluorescence spectroscopy

To check the effect of EDTA on the thermal stability of PduMCP, the purified PduMCP samples are incubated in the absence and presence of EDTA at different concentrations for 5 min at room temperature. Temperature dependent intrinsic fluorescence assay for the EDTA treated and control samples of PduMCP is carried out using Spectrofluorimeter FS5 (Edinburgh Instruments, UK). The sample are excited at 280 nm (band width = 1 nm) and emission spectra (band width = 1 nm) are recoded between 300 to 500 nm at temperatures between 20°C-90°C. The 1^st^ derivative of the emission maxima versus temperature plot reveals the melting temperatures (*T_m_* values) of PduMCP.

### Transmission electron microscopy

TEM imaging of the shell protein PduBB’ is perfromed by drop casting 10 μl of 0.2 mg/ml of the shell protein on to carbon coated TEM grid. This is followed by staining with 7 μl of 1 % (w/v) uranyl nitrate (freshly prepared and filtered). The grid is then washed with Milli-Q and air dried for 24 h at room temperature in dark. TEM image is obtained using JEM 2100 TEM (JEOL, USA) operated at 120 kV.

## Acknowledgements

This work is supported by SERB, India Grant EMR/2015/000746 and DBT grant BT/12/IYBA/2019/08 to S.S. G.K thanks INST, Mohali for fellowship.

## Author Contributions

SS conceived the idea. GK performed the experiments. GK and SS wrote the manuscript.

## Supporting Figures

**Figure S1:**
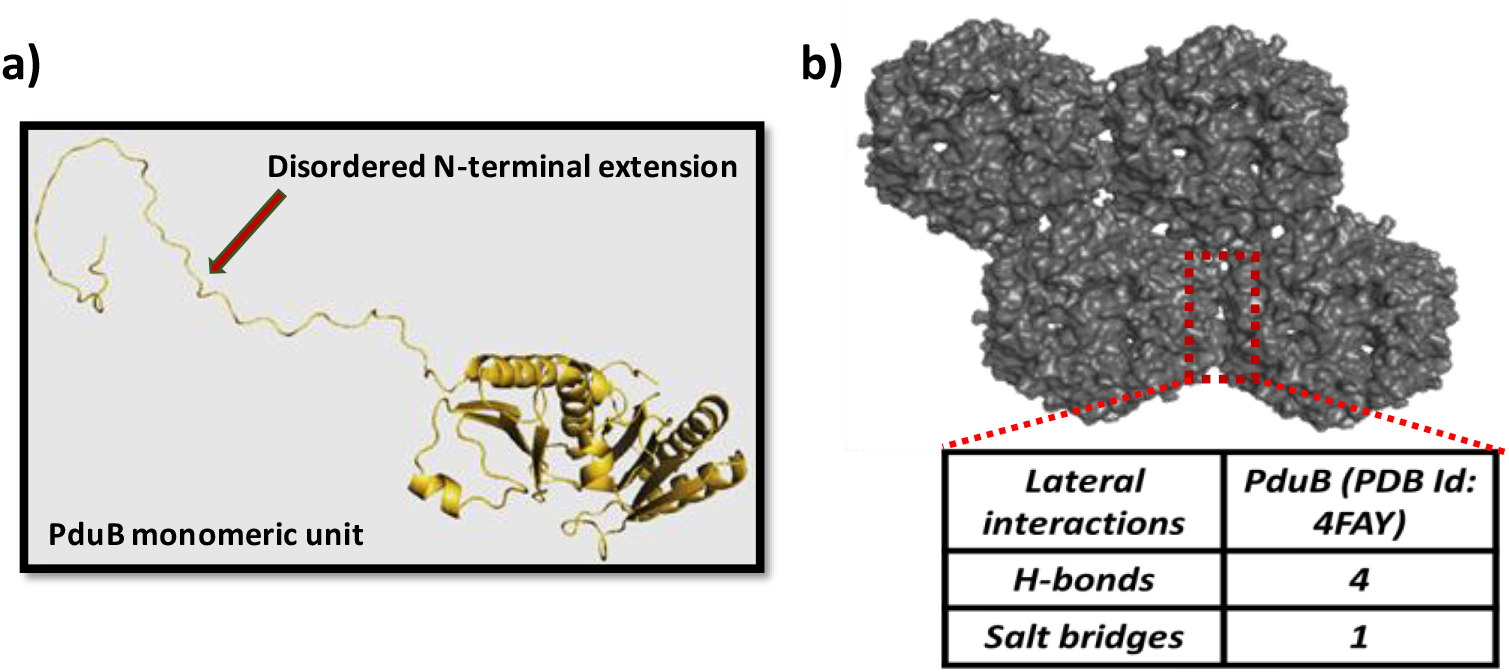
a) *Ab-initio* modeling of monomeric unit of PduB (270 amino acids using AIDA server, b) Edge-Edge lateral association in shell protein PduB from *Lactobacillus reuteri* (PDB ID: 4FAY)

**Figure S2:**
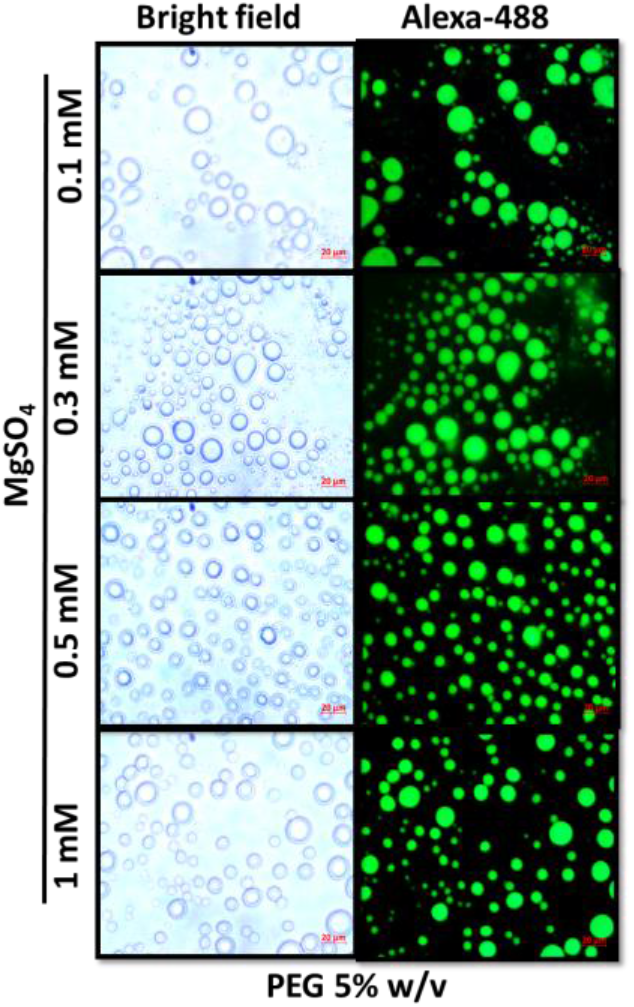
Liquid-liquid phase separation of PduBB’ at MgSO_4_ concentrations between 0.1 mM to 1mM.

**Figure S3:**
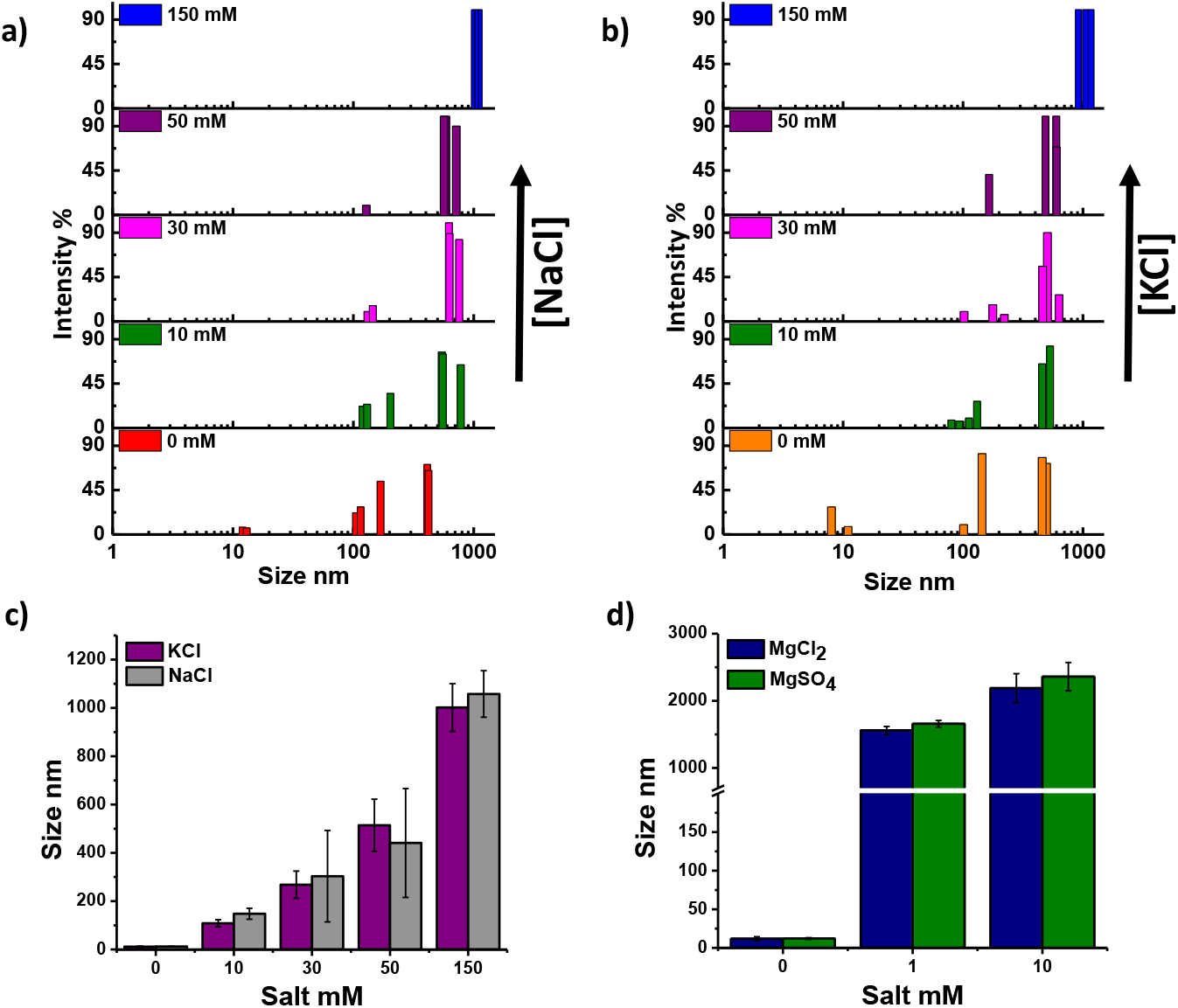
a) Size distribution of shell protein PduBB’ under NaCl concentrations varied from 0 mM to 150 mM, b) Size distribution of shell protein PduBB’ under KCl concentrations varied from 0 mM to 150 mM, c) Size of PduBB’ assemblies in solution phase in the presence of KCl and NaCl at concentrations ranging from 0 mM to 150 mM, d) Size of PduBB’ assemblies in solution phase in the absence and presence of presence of MgCl_2_ and MgSO_4_ (1mM and 10 mM).

### Discussion

#### Figure S3: Salt promotes self-association of the shell protein PduBB’

To understand how change in ionic strength of environment affects the PduBB’ assemblies, we look at the size distribution of PduBB’ (0.1 mg/ml) in solution phase without any crowding agent, in the absence and presence of NaCl (10 mM-150 mM) using dynamic light scattering (DLS) (Fig. S5a). In solution, PduBB’ (0.1 mg/ml) shows a wide size distribution of shell assemblies (intensity percentage in DLS) between 10nm-500nm. This suggests that a heterogeneous population of shell protein assemblies exists in solution phase. Increase in NaCl concentration results in self-association of PduBB’ in solution. At 50 mM NaCl, more than 90% of the size distribution is seen above 500 nm. Increase in NaCl concentration to 150 mM, results in further self-association of the shell protein with a majority of PduBB’ assemblies falling in the micron range. A similar trend is also noticed when PduBB’ is incubated in the presence of KCl (Fig. S5b). The self-association behavior of PduBB’ in the presence of NaCl or KCl becomes more apparent when we look at the maximum population (number percentage in DLS) of the shell protein assemblies in solution at different ionic strengths (Fig. S5c). In the absence of NaCl or KCl, the solution containing PduBB’ shows a maximum population of shell protein assemblies around 10 nm which changes to a maximum population in the micron range upon increasing the salt concentration to 150 mM. Interestingly, the effect of Mg^2+^ salts on PduBB’ assembly is found to be more aggressive than the monovalent salts (NaCl and KCl), with PduBB’ showing maximum population of size above 1000 nm at 1 mM and above 2000 nm at 10 mM of MgCl_2_ or MgSO_4_ (Fig. S5d). Our data indicate that divalent salts (MgCl_2_ and MgSO_4_) have a higher potential than monovalent salts (NaCl and KCl) to promote self-association of PduBB’ in solution.

**Figure S4:**
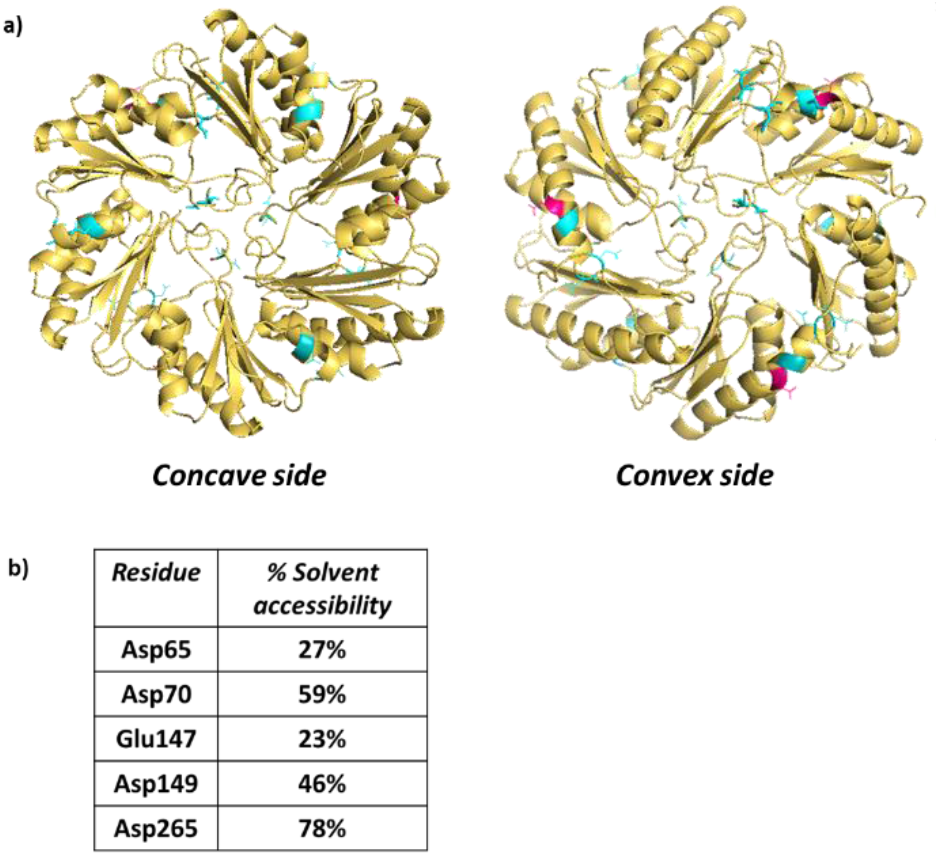
a) Docking of PduBB’ with Mg2+ ions using MIB webserver. The Asp and Glu residues predicted to bind Mg^2+^ ions are shown in cyan and pink respectively, b) Percentage solvent accessibility of the Asp/Glu residues predicted to bind to Mg^2+^ ions.

### Discussion

#### Figure S4: Docking of PduBB’ with Mg2+

MIB webserver is programmed to identify the divalent cation binding sites on proteins theoretically predicted the residues Asp65, Asp70, Glu147, Asp149, and Asp265 in PduBB’ that have the potential to bind Mg^2+^ ions **(Fig. S4a)**. Our bioinformatics analysis suggests that these residues in each chain of PduBB’ are considerably exposed to solvent, thereby, increasing their probability of binding to Mg^2+^ ions **(Fig. S4b)**.

**Figure S5:**
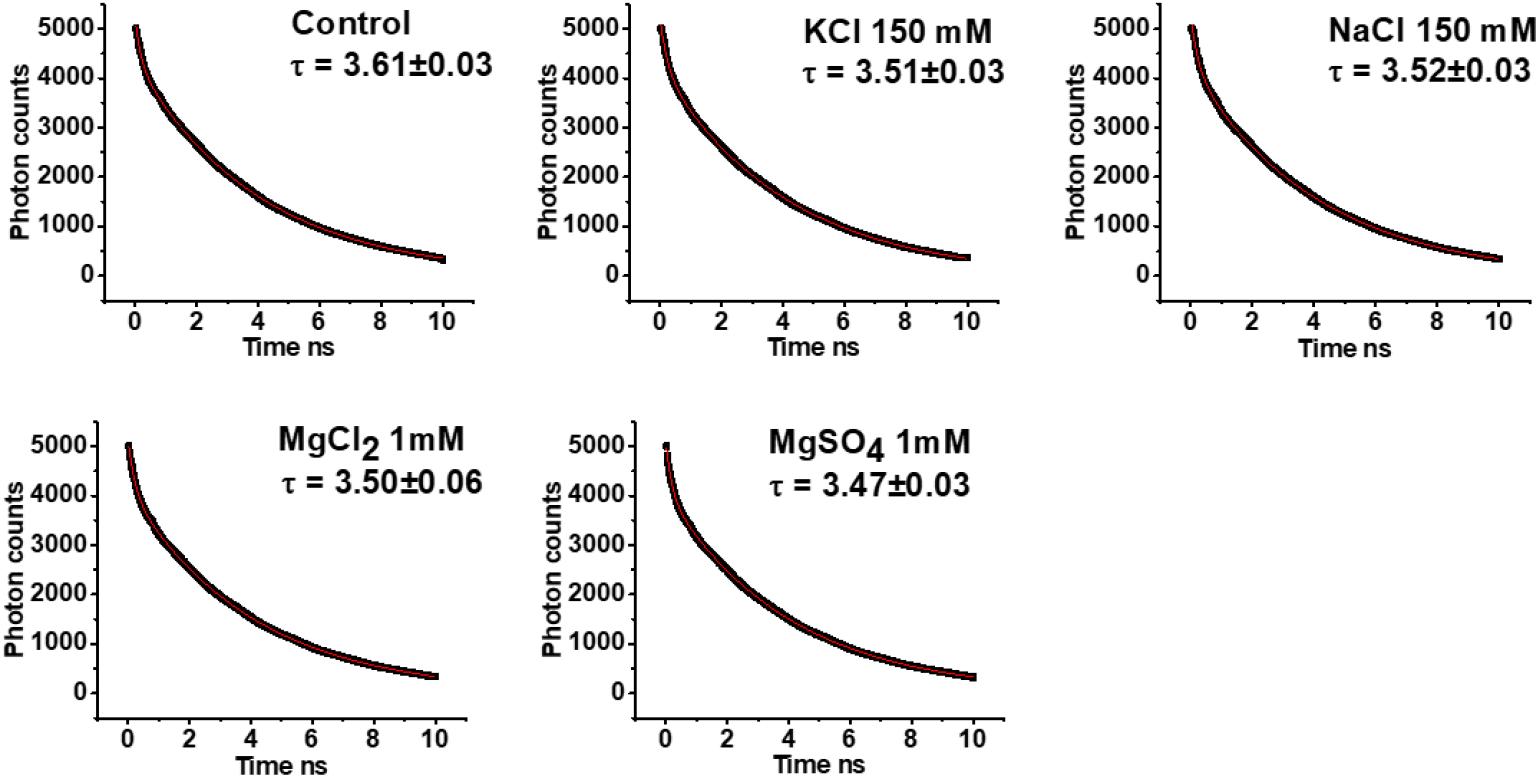
Fluorescence lifetime decay of Alexa-488 labeled PduBB’ (a) and in the presence of salts, KCl 150 mM (b), NaCl 150 mM (c), MgCl_2_ 1 mM (d), MgSO_4_ 1 mM (e). The lifetime decay curve was fitted to monoexponential decay equation after converting the photon count axis into linear scale.

**Figure S6:**
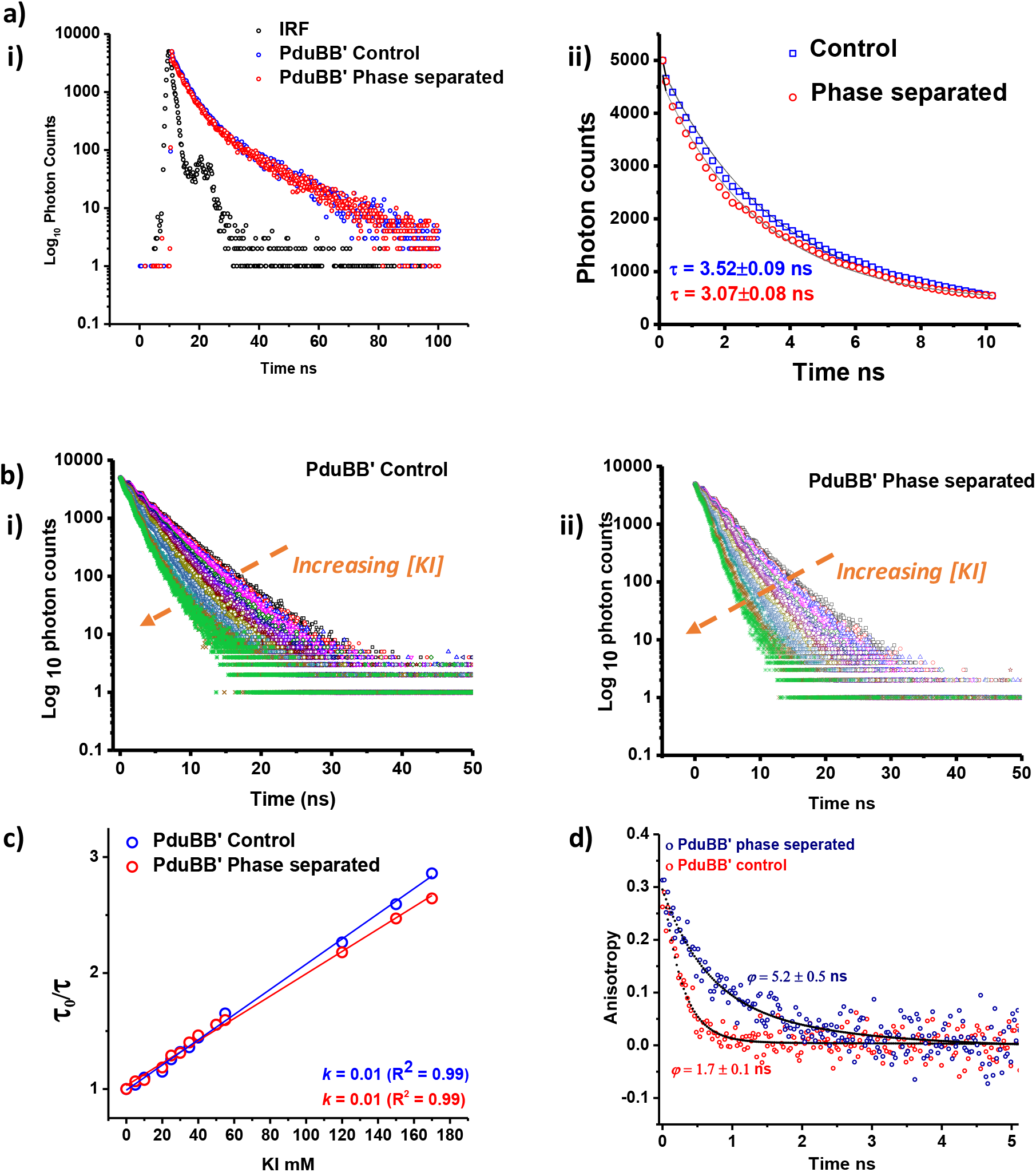
a) Fluorescence lifetime of Alexa-488 labeled PduBB’ (0.5 mg/ml) under (i) control and (ii) phase separated condition (1mM MgSO_4_ and 5% w/v PEG-6000), b) Fluorescence lifetime of PduBB’ under (i) control and (ii) phase separated condition (1mM MgSO_4_ and 5% w/v PEG-6000) in the presence of increasing concentrations of potassium iodide (KI), c) Stern-Volmer plot of τ_0_/ *τ* (τ_0_ = fluorescence lifetime of Alexa-488 labeled PduBB’ at 0 mM KI, τ = fluorescence lifetime of Alexa-488 labeled PduBB’ at different concentrations of KI, d) Time resolved fluorescence anisotropy of PduBB’ under control environment and post phase separation.

### Discussion

#### Figure S6: Probing the local environment of PduBB’ during LLPS

In this section we discuss how phase separation affects the local environment around shell protein PduBB’. We determine fluorescence lifetime of Alexa-488 labelled PduBB’ under control and phase separated condition (1 mM MgSO_4_, 5% w/v PEG-6000) (Fig. S3a, i). Fitting the first 10 ns seconds of lifetime decay curve using monoexponential decay equation, shows a small decrease in lifetime post phase separation by ~0.5 ns (Fig. S3a, ii). This change in lifetime may be due to the alteration in the microenvironment of fluorophore molecule due to local enrichment of ions which tend to mask the surface charge of the shell protein during phase separation. We next perform solvent accessibility of Alexa-488 labelled PduBB’ under control and phase separated condition, using water soluble quencher potassium iodide (KI) (Fig. S3b, i and ii). In both cases the fluorescence lifetime of labelled PduBB’ decreases with the increase in KI concentration. A plot of τ_0_/τ at different concentrations of KI (Stern-Volmer plot) where τ_0_ is the fluorescence lifetime of Alexa-488 labeled PduBB’ at 0 mM KI and τ is the fluorescence lifetime of Alexa-488 labeled PduBB’ at different concentrations of KI, is linear in both solution and phase separated condition (Fig. S3c). Fitting the Stern-Volmer plots using linear fit equation shows same binding constants for the quencher in case of both control and phase separated PduBB’ sample. This suggests that the accessibility of KI to the shell protein backbone is similar in case of both control and phase separated samples. In other words, phase separation does not result in disruption of the solvation layer of the shell protein backbone. The solvation of peptide backbone of PduBB’ may be the reason why the shell protein behaves like liquid post phase separation and not collapse into aggregates. Further, in time resolved anisotropy studies we observed a rotational correlation time of 1.7±0.1 ns in solution, and of 5.2±0.5 ns in LLPS PduBB’ (Fig. S3d). The higher rotational correlation time, suggests a restricted rotational dynamics of the shell protein in the crowded micro-environment. Together our fluorescence lifetime and anisotropy studies indicate that during the LLPS process the shell protein backbone remains solvated but have restricted rotational dynamics.

**Figure S7:**
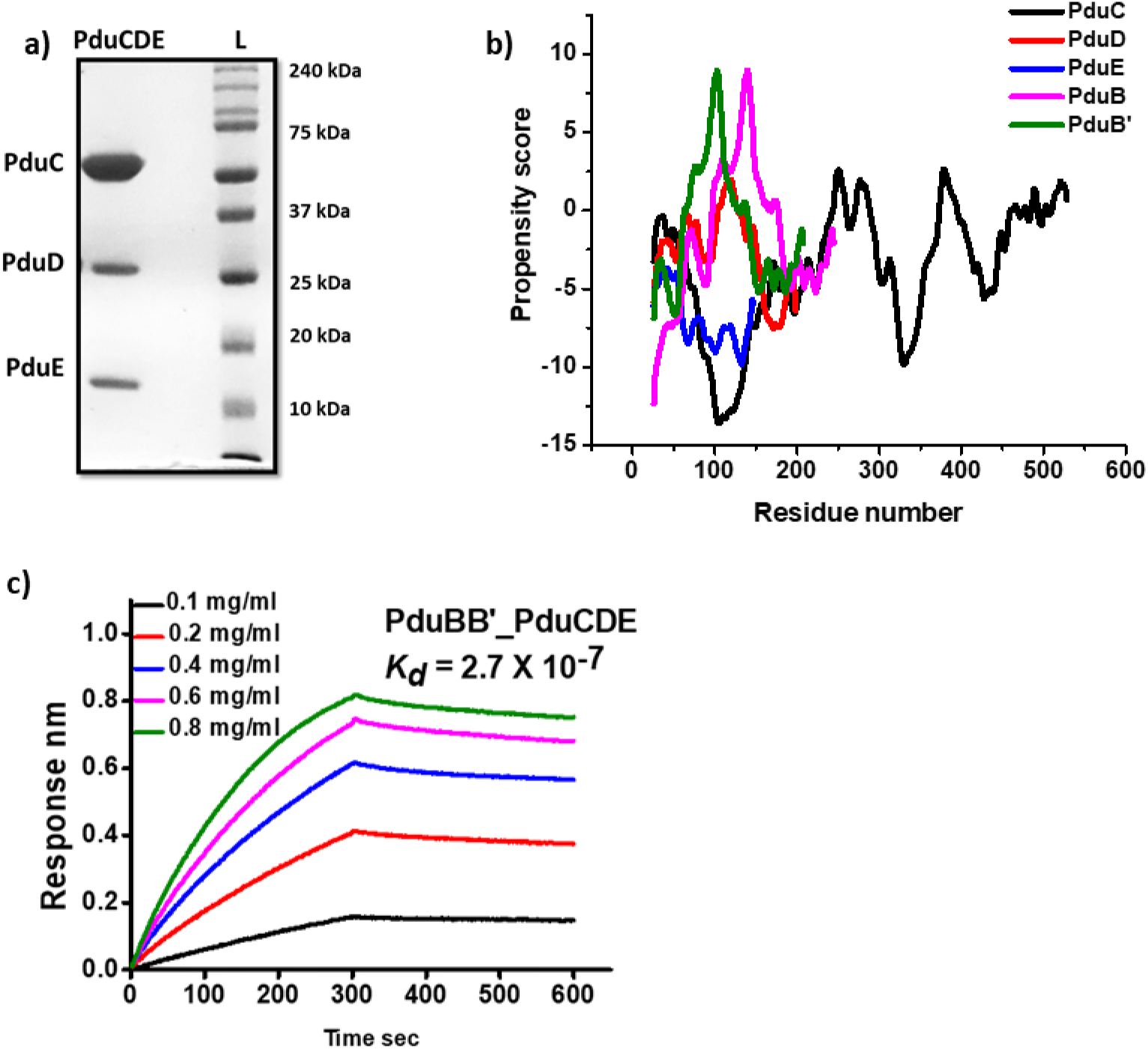
a) SDS PAGE of PduCDE, b) LLPS propensity scores of the three subunits of PduCDE and the monomeric units of shell protein PduBB’, c) Biolayer interferometry to probe the affinity of PduBB’ towards PduCDE.

**Figure S8:**
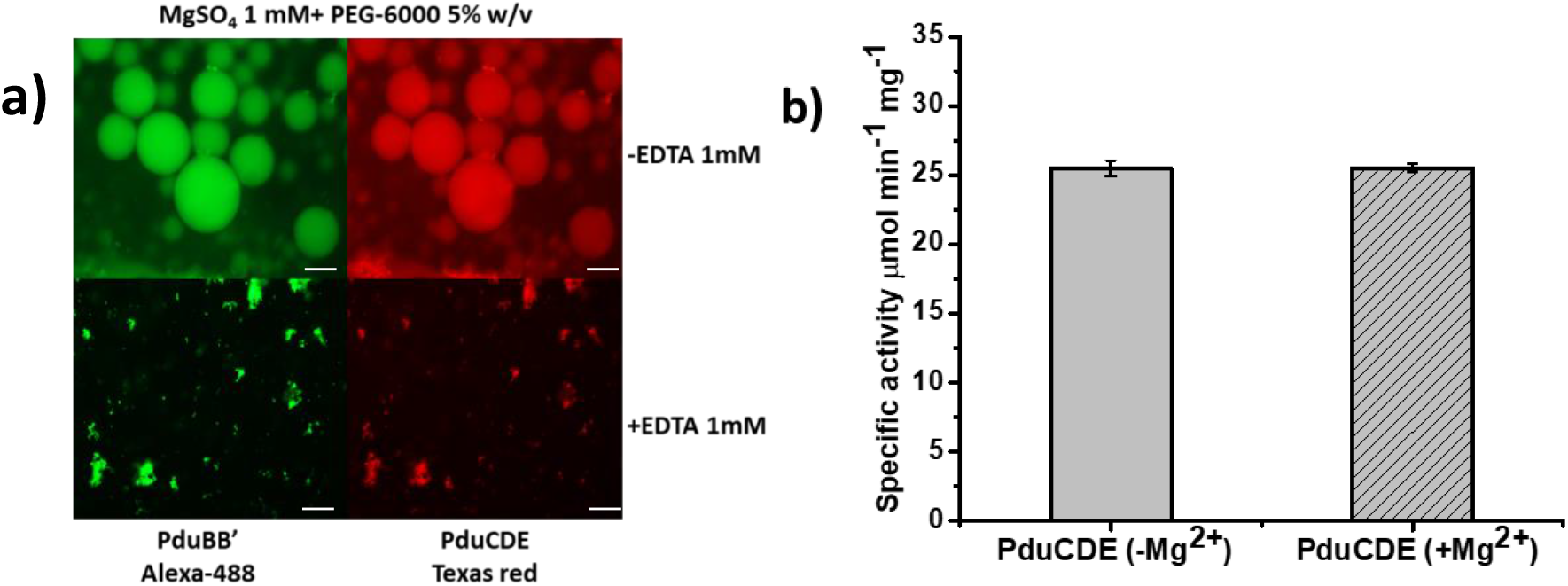
a) Phase separation of PduBB’ and PduCDE mixture in the absence and presence of 1 mM EDTA, scale bar = 10 μm, b) Specific activity of PduCDE in the absence and presence of 1 mM MgSO_4_.

**Figure S9:**
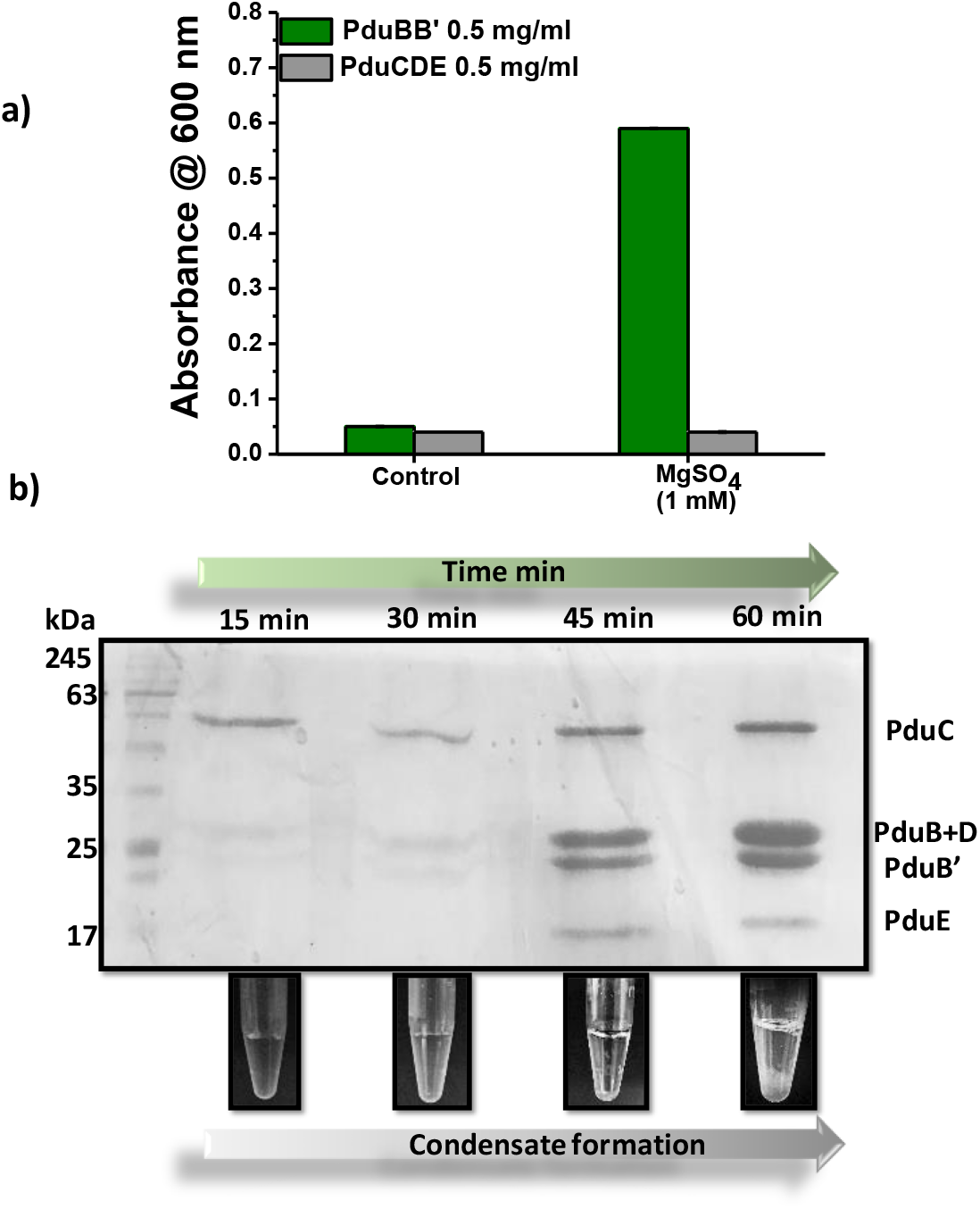
a) Turbidity assay of PduBB’ (0.5 mg/ml) and PduCDE (0.5 mg/ml) in the absence and presence of 1 mM MgSO_4_, b) Time dependant Co-condensation of PduBB’ and PduCDE in solution.

### Discussion, Figure S9

a) The effect of 1 mM MgSO_4_ on solutions of PduBB’ and PduCDE is checked by turbidity assay. In the absence of the salt, both PduBB’ and PduCDE (0.5 mg/ml each) gives clear solution without any turbidity. In the presence of 1 mM MgSO_4_ the solution of PduBB’ turns turbid in an hour giving high optical density, while PduCDE solution remains clear. This suggests that shell protein PduBB’ has a higher potential to undergo self-association in solution than the enzyme PduCDE. b) We check if the mixture of shell protein and enzyme has the ability to co-separate out in solution. For this study, we mix the shell protein PduBB’ and enzyme PduCDE (0.5 mg/ml each in final mixture) in the presence of 1 mM MgSO_4_, and condensate formed at different time interval is run on SDS-PAGE gel. We observe that, with time the intensity of the bands corresponding to PduBB’ and PduCDE inccreases on gel. This indicates that in the presence of 1 mM MgSo4 triggers the self-assembly of PduCDE and PduBB’ in solution phase.

**Figure S10:**
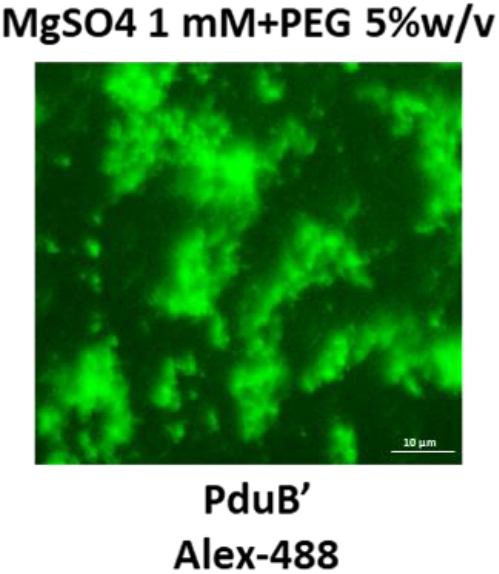
Solid associates of PduB’ (233 amino acids) seen under phase separated condition.

## Supporting Video

**Video Sv 1: Fusion events for the shell protein PduBB’ (1 mg/ml, in the presence of 5% w/v PEG-6000 and 1 mM MgSO_4_), as seen under fluorescence microscope**.

